# Why architecture matters: Controlling gene expression through design

**DOI:** 10.64898/2026.08.06.743186

**Authors:** Abhilasha Gupta, Mitchell Lewis

## Abstract

Inducible gene expression systems are widely used in synthetic biology and gene therapy, yet their performance depends not only on regulator chemistry but also on circuit architecture. Here, we examine how promoter organization shapes TetR-based gene regulation in mammalian cells using a panel of single-vector constructs spanning a broad range of promoter strengths.

Experiments and thermodynamic models show that bidirectional circuits impose a trade-off between output and control: increasing promoter strength elevates both induced and basal expression, compressing dynamic range. Incorporating transcriptional coupling explains the parallel scaling of these states in compact divergent designs. In contrast, autogenous regulation couples repressor production to transcription, introducing negative feedback that buffers promoter strength and preserves fold induction.

Finally, adding ligand-responsive aptazymes as a post-transcriptional layer further suppresses basal expression while maintaining inducibility, albeit with reduced maximal output. Together, these results identify regulatory architecture as a primary determinant of circuit performance and establish design principles for constructing more predictable gene expression systems in eukaryotic cells.

## Introduction

Precise control of gene expression is central to both basic studies of gene regulation and the development of therapeutic gene delivery systems. The conceptual foundation for this work traces to the operon model, in which regulatory proteins control transcription by binding specific DNA operators, and to allosteric theory, in which ligand binding shifts the conformational equilibrium of a regulatory protein [1, 2]. These principles provide the basis for small-molecule-controlled gene circuits: ligand binding changes regulator activity, regulator occupancy changes promoter state, and promoter state determines transcriptional output.

Tetracycline-responsive systems are among the most widely used examples of this design logic in mammalian cells. The original Tet-Off system converted the bacterial Tet repressor into a eukaryotic transcriptional activator by fusing TetR to VP16, allowing tetracycline or doxycycline to turn expression off by preventing DNA binding [3]. The subsequent Tet-On system reversed regulatory polarity by engineering a reverse Tet transactivator that activates transcription in the presence of tetracycline-class ligands [4]. Further engineering expanded the sequence space of tetracycline-dependent transactivators and improved sensitivity, basal activity, and dynamic range [5]. Together, these systems established Tet-based control as a modular and broadly useful platform for reversible, small-molecule-regulated gene expression.

The success of Tet systems reflects several practical advantages: TetR and tetO are orthogonal to most mammalian regulatory networks, the input ligand is cell-permeable and experimentally convenient, and the system can be configured either as ligand-repressed expression (Tet-Off) or ligand-induced expression (Tet-On) [3, 4, 6]. However, the regulatory behavior of these systems is not determined by ligand affinity alone. TetR function depends on allosteric coupling between ligand binding and operator binding, such that doxycycline shifts the population of TetR away from the DNA-binding-competent state [7]. As a result, circuit performance depends on both the molecular free-energy landscape of the regulator and the architecture in which that regulator is embedded.

Although much effort has focused on engineering regulatory proteins, activation domains, ligand-binding properties, and operator sequences, the overall performance of an inducible system also depends on how its components are arranged within a genetic circuit. In particular, the organization of promoters, operators, and coding sequences—referred to here as the *regulatory architecture*—plays a critical role in determining basal expression, induced output, dynamic range, and robustness. Thermodynamic models of transcription provide a natural framework for analyzing these effects because expression can be written in terms of promoter-state probabilities and statistical weights for RNA polymerase, repressor, and ligand-dependent regulatory states [8–10]. Thus, different architectures can produce qualitatively distinct behaviors even when built from the same molecular components.

Early implementations of inducible gene expression systems in mammalian cells typically used a two-vector architecture in which the regulatory protein and target gene were encoded on separate constructs [3, 4]. In this design, one vector constitutively expresses the transcription factor, whereas a second vector contains a regulated promoter controlling the gene of interest. This separation allows the two components to be tuned independently: regulator abundance can be adjusted through promoter strength or plasmid ratio, while target gene output is determined by the regulated promoter. However, this flexibility comes at the cost of cell-to-cell variability, because the two vectors are delivered and maintained independently.

To reduce this variability, many designs have shifted toward single-vector or “all-in-one” architectures in which both the regulatory protein and the target gene are encoded on the same construct. By physically linking these components, single-vector systems help ensure that each cell receives a complete regulatory module, improving stoichiometric consistency relative to two-vector delivery. This feature is especially important in viral gene delivery, where vector capacity, copy number, and co-delivery efficiency impose practical constraints [21]. At the same time, placing multiple transcriptional units in close proximity can introduce new interactions, including promoter–promoter coupling, transcriptional interference, RNA polymerase traffic, and local chromatin effects [11].

Within single-vector systems, one compact strategy uses *bidirectional* or divergent promoters, in which one promoter drives expression of the regulatory protein and an oppositely oriented promoter controls the target gene. In this configuration, repressor production and target expression are nominally independent but physically linked. This arrangement is attractive for compact vector design, but it can create functional coupling between the two promoters such that changes in one promoter alter the activity of the other.

An alternative strategy uses *autogenous* regulation, in which the regulatory protein is expressed from the same promoter that it controls. In this architecture, repressor production is directly coupled to transcriptional output, creating a negative-feedback loop. Increased promoter activity produces more repressor, which suppresses further transcription. Negative feedback can stabilize expression, reduce noise, and buffer parameter variation, but it can also constrain maximal output and alter the relationship between promoter strength and dynamic range [12–14, 20].

A third class of architectures introduces *layered regulation* by combining transcriptional control with post-transcriptional mechanisms. Ligand-responsive RNA elements such as aptazymes can be inserted into transcripts to modulate mRNA stability or translation in a ligand-dependent manner. When combined with transcriptional repression, these systems provide multiple points of control, enabling stronger suppression of basal expression and improved regulatory stringency, albeit often at the cost of reduced maximal output [15–17].

Despite the widespread use of Tet-based regulatory systems, a quantitative understanding of how circuit organization shapes regulatory behavior remains incomplete. In particular, it is not sufficient to ask whether a regulator is inducible or co-repressible, or whether a ligand binds tightly. Instead, one must ask how promoter strength, regulator abundance, ligand-dependent operator occupancy, and promoter–promoter interactions are integrated within a given architecture to determine basal expression, induced output, and dynamic range.

## Results

Here, we examine how regulatory architecture shapes the behavior of TetR-based inducible systems. By combining experiment with thermodynamic modeling, we can define the constraints imposed by different architectural designs and establish a rational framework for selecting among them, particularly in the context of regulating gene expression in mammalian cells.

### Bidirectional architecture: Promoter coupling and regulatory consequences

Bidirectional regulatory systems provide a compact strategy for coordinating expression of a regulatory protein and its target gene within a single genetic construct. By positioning two promoters in opposing orientations on the same DNA segment, these systems enable simultaneous expression of the repressor and the regulated transgene while minimizing vector size. This organization is particularly advantageous for applications such as viral gene delivery, where cargo capacity is limited and co-delivery of multiple vectors is inefficient. In addition, single-vector bidirectional systems ensure that each transduced cell receives both regulatory components, improving stoichiometric consistency relative to multi-vector approaches.

In the architecture used here, one promoter (*P_R_*) drives constitutive expression of TetR, whereas a second promoter (*P_G_*), oriented in the opposite direction, controls expression of a luciferase reporter gene. The two promoters are separated by an intervening transcriptional pause/insulator sequence designed to reduce transcriptional interference and limit promoter cross-talk between the divergent transcription units [11, 21]. A tetracycline operator sequence positioned downstream of *P_G_*permits TetR binding and repression of reporter transcription. Addition of doxycycline reduces TetR DNA-binding affinity, causing TetR dissociation from the operator and relieving repression, thereby allowing transcriptional activation. Because repressor production and reporter expression are driven by separate promoters, the system initially appears to permit independent tuning of regulatory strength and transcriptional output.

Despite this apparent independence, it is possible that the close physical proximity of the two promoters creates the potential for functional coupling. Transcriptional initiation at one promoter can influence activity at the opposing promoter through mechanisms such as transcriptional interference, local changes in DNA topology, or cooperative stabilization of transcriptional complexes. Although biidirectional systems provide a compact regulatory design, they also create a framework in which promoter–promoter interactions can shape circuit behavior.

To systematically examine how promoter strength influences regulatory performance, we generated a panel of bidirectional TetR modules in which the strengths of *P_R_*and *P_G_*were varied independently across a broad transcriptional range. Promoters were selected to represent weak, intermediate, and strong expression regimes based on prior characterization in mammalian cells (Table 1). Weak promoters, such as hPGK and CMV_min_, provide low basal transcription and are less susceptible to saturation effects but may limit maximal output. Intermediate promoters, including EF1*α* variants and CMV–IE, provide a balance between expression strength and regulatory control. Strong promoters, such as CMV and CAG, maximize transcriptional output but can overwhelm repression, resulting in elevated basal expression.

**Table 1.** Promoter strengths used in the bidirectional regulatory constructs.

| Category | Promoter | RLU | Relative |
| --- | --- | --- | --- |
| Strong | CMV | $2.9 \times 10^5$ | 97 |
| Medium | CAG | $1.5 \times 10^5$ | 50 |
| | CMV–IE | $1.5 \times 10^5$ | 50 |
| | EF1α (long) | $9.0 \times 10^4$ | 30 |
| Weak | EF1α (core) | $2.5 \times 10^4$ | 8.3 |
| | CMV<sub>min</sub> | $5.0 \times 10^3$ | 1.7 |
| | hPGK | $3.0 \times 10^3$ | 1 |

Relative luminescence units (RLU) report luciferase activity and are normalized to the weakest promoter (hPGK).

All pairwise promoter combinations were assembled into single-vector bidirectional constructs, generating a matrix in which repressor production and reporter transcriptional capacity could be varied systematically. Constructs were transiently transfected into mammalian cells under identical conditions, and reporter expression was measured in both the absence and presence of saturating doxycycline (Fig. 1). This design enabled direct comparison of basal expression, induced output, and fold induction across promoter combinations.

**Fig 1.**
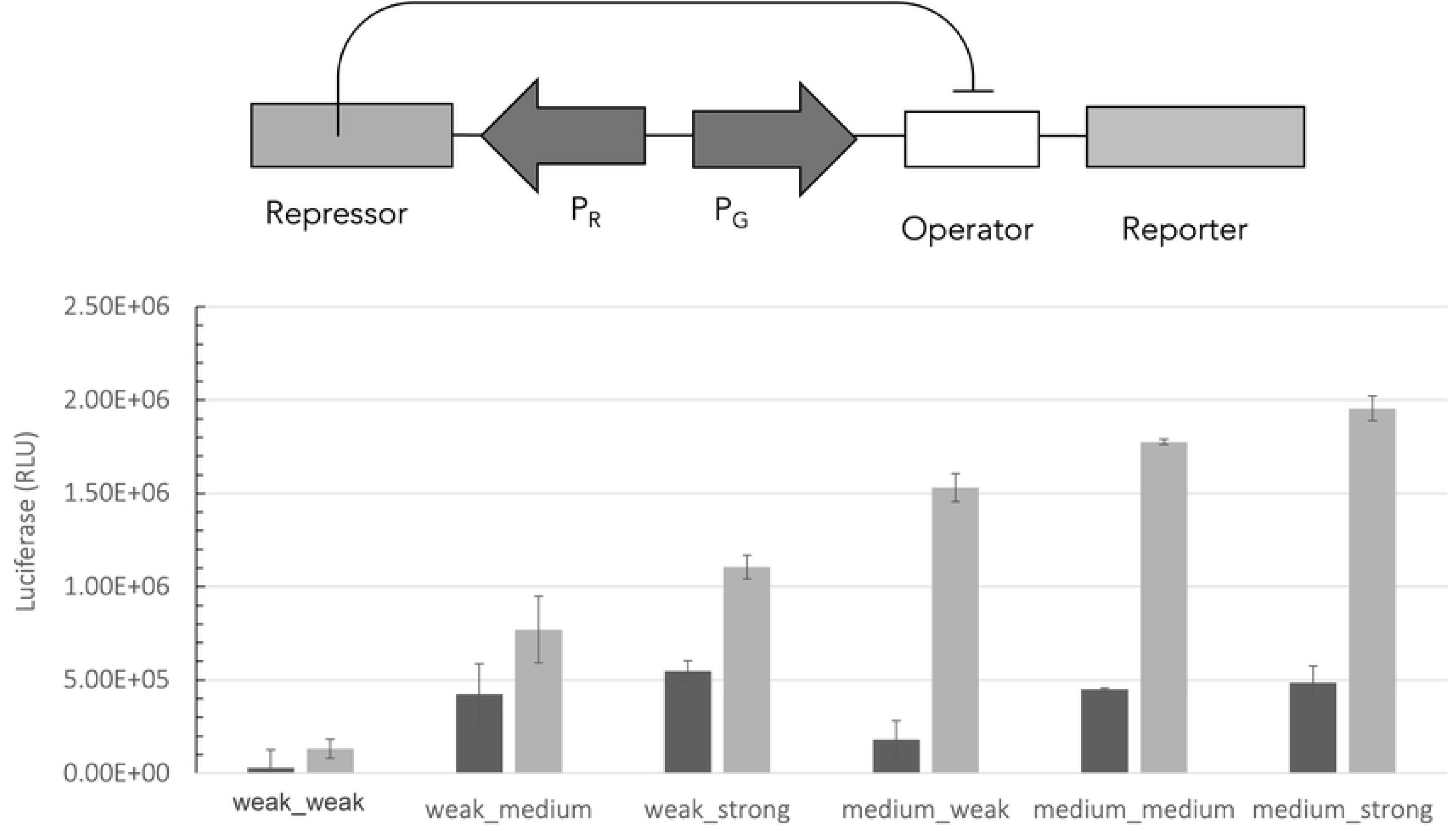
Bidirectional TetR regulatory architecture and promoter-dependent reporter output. Opposing promoters flank a shared Tet operator such that TetR expressed from one promoter regulates transcription from the other. Reporter activity is shown for combinations of repressor and reporter promoter strengths. Dark bars denote basal expression; light bars denote induced expression (+doxycycline). Error bars represent mean ± SD (*n* = 8). Reporter promoter strength primarily determines output amplitude, whereas repressor-side promoter strength also alters induced expression, consistent with functional promoter coupling.

Reporter expression varied systematically with the strengths of both promoters. Increasing the strength of the reporter promoter (*P_G_*) produced a monotonic increase in both basal and induced expression, consistent with increased RNA polymerase occupancy at the regulated promoter. Thus, *P_G_* primarily determines the overall transcriptional capacity of the circuit.

Unexpectedly, increasing the strength of the repressor promoter (*P_R_*) also increased reporter expression in both the basal and induced states. Rather than selectively reducing basal expression through increased TetR abundance, stronger repressor promoters elevated overall transcriptional output. Basal and induced expression therefore scaled together across promoter combinations, indicating that the two promoters do not function independently within the bidirectional architecture.

This coordinated scaling compressed the achievable dynamic range at higher promoter strengths. Although stronger promoters increased absolute induced expression, they also elevated basal expression, limiting separation between OFF and ON states. A modest reduction in induced expression with increasing repressor strength further suggests that repression is not completely relieved even in the presence of doxycycline, consistent with residual operator occupancy.

These observations are inconsistent with a strictly independent promoter model. If the two promoters acted independently, increasing *P_R_* strength would primarily increase repressor abundance and reduce basal reporter expression while leaving the fully induced state largely unchanged. Instead, induced reporter expression also increased with *P_R_* strength, indicating that transcriptional activity at one promoter influences output from the opposing promoter. The data suggest that promoter activity is functionally interconnected.

### A thermodynamic model reveals promoter coupling

To interpret the behavior of the bidirectional regulatory system, we developed a statistical thermodynamic model in which gene expression is proportional to the probability that RNA polymerase (RNAP) occupies a given promoter [8–10]. In this framework, transcription is determined by the equilibrium distribution of promoter occupancy states, with RNAP binding and repressor interactions explicitly enumerated.

The system comprises two promoters: a repressor promoter (*P_R_*), which drives expression of the regulatory protein, and a reporter promoter (*P_G_*), which is subject to repression. RNAP binding to each promoter is described by the dimensionless weights

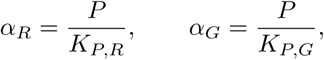

where *P* is the effective RNAP concentration and *K_P,R_*and *K_P,G_*are the corresponding dissociation constants. Repressor binding is described by

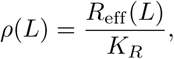

where *R*_eff_ (*L*) = *p_A_*(*L*) *R*_tot_ is the ligand-dependent concentration of DNA-binding–competent repressor.

All allowed promoter occupancy states are enumerated and assigned statistical weights based on RNAP and repressor binding. The repressor blocks RNAP binding at the reporter promoter but does not interfere with binding at the repressor promoter. Interactions between RNAP molecules at the two promoters are captured by a coupling parameter *ω*, which modifies the statistical weight of states in which both promoters are simultaneously occupied. The allowed promoter states and their statistical weights are summarized in Table 2.

**Table 2.** Promoter occupancy states and corresponding statistical weights.

| State | Weight |
| --- | --- |
| Empty DNA | 1 |
| RNAP bound at $P_R$ | $\alpha_R$ |
| RNAP bound at $P_G$ | $\alpha_G$ |
| RNAP bound at both promoters | $\omega \alpha_R \alpha_G$ |
| Repressor bound | $\rho$ |
| Repressor + RNAP at $P_R$ | $\rho \alpha_R$ |

### Independent promoter model

As a baseline, we first consider the uncoupled limit (*ω* = 1), in which the two promoters behave independently. In this case, reporter expression is determined solely by occupancy of the reporter promoter,

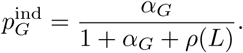

Because the promoters are independent, reporter expression depends only on *α_G_*and *ρ*(*L*) and is therefore insensitive to the strength of the repressor promoter (*α_R_*) except through its effect on total repressor abundance. Increasing *α_R_* is thus predicted to reduce basal expression while leaving induced expression largely unchanged.

Experimentally, however, increasing the strength of the repressor promoter leads to increases in both basal and induced expression. This behavior is inconsistent with the independent promoter model, indicating that the two promoters are functionally coupled.

### Coupled-promoter model

In this context, coupling refers to a functional interaction in which transcriptional activity at one promoter influences transcription from the opposing promoter, despite each promoter having its own regulatory elements. As a result, increasing transcriptional activity at either promoter can enhance activity from the opposing promoter, causing basal and induced expression to scale together rather than independently. To account for this behavior, we consider the full model with *ω ≠* 1. The partition function is

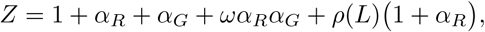

and reporter expression is given by

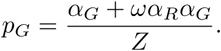

The coupling parameter *ω* quantifies promoter–promoter interactions. For *ω >* 1, simultaneous RNAP occupancy of both promoters is favored, effectively linking repressor production and reporter transcription.

In this coupled regime, increasing *α_R_* enhances not only repressor production but also reporter transcription through the *ωα_R_α_G_* term. This produces coordinated increases in both basal and induced expression, consistent with the observed parallel scaling across promoter combinations.

Global fits to the full dataset yield coupling strengths in the range *ω* ∼ 2–5, corresponding to a 2–5-fold enhancement of joint promoter occupancy. The coupled model quantitatively reproduces the observed expression profiles, whereas the independent model fails systematically. This improvement is reflected in goodness-of-fit metrics, with the independent model yielding *R*^2^ ≲ 0.6 and the coupled model achieving *R*^2^ ≳ 0.85.

These results demonstrate that promoter–promoter interactions are both necessary and sufficient to explain the behavior of bidirectional regulatory circuits. In contrast to the independent model, in which repression and transcriptional capacity can be tuned separately, the coupled-promoter architecture intrinsically links these processes. As a result, promoter strength acts as a shared determinant of transcriptional output in both basal and induced states, imposing a fundamental constraint on achievable dynamic range.

### Autogenous architecture: Feedback and regulatory behavior

An alternative strategy for inducible gene regulation is to completely couple the repressor and transgene using an autogenous architecture, in which the regulatory protein is expressed from the same promoter that it regulates. This configuration creates a built-in negative-feedback loop that directly couples repressor production to transcriptional output. Negative feedback is widely used in natural and synthetic gene networks because it stabilizes gene expression, reduces sensitivity to fluctuations in promoter activity, and improves robustness across heterogeneous cellular environments [12–14, 18, 20]. These properties are particularly advantageous in compact single-vector systems, where independent tuning of regulatory components is limited and stable circuit behavior must emerge from the architecture itself.

In the autogenous architecture used here, a single promoter (*P*) drives expression of both TetR and a downstream luciferase reporter. The two proteins are encoded within the same transcriptional unit and separated by a self-cleaving 2A peptide, allowing coordinated expression while producing independent protein products [24, 25]. A tetracycline operator positioned downstream of the promoter enables TetR to repress transcription from the same promoter that drives its own expression. In the absence of doxycycline, TetR binds the operator and suppresses transcription, thereby reducing production of both the repressor and the reporter. Addition of doxycycline decreases TetR DNA-binding affinity, relieving repression and increasing transcription [3, 4, 6, 26]. Because repressor abundance depends directly on promoter activity, increased transcription automatically increases repression, establishing a self-limiting negative-feedback loop.

We created a series of autogenous modules in which the intrinsic strength of the shared promoter was varied across the same panel of weak, intermediate, and strong promoters used in the bidirectional system. Constructs were transiently transfected into mammalian cells under identical conditions, and reporter expression was quantified by luciferase assay in both the absence and presence of saturating doxycycline. These measurements enabled direct comparison of basal expression, induced output, and fold induction across promoter strengths (Fig. 2).

**Fig 2.**
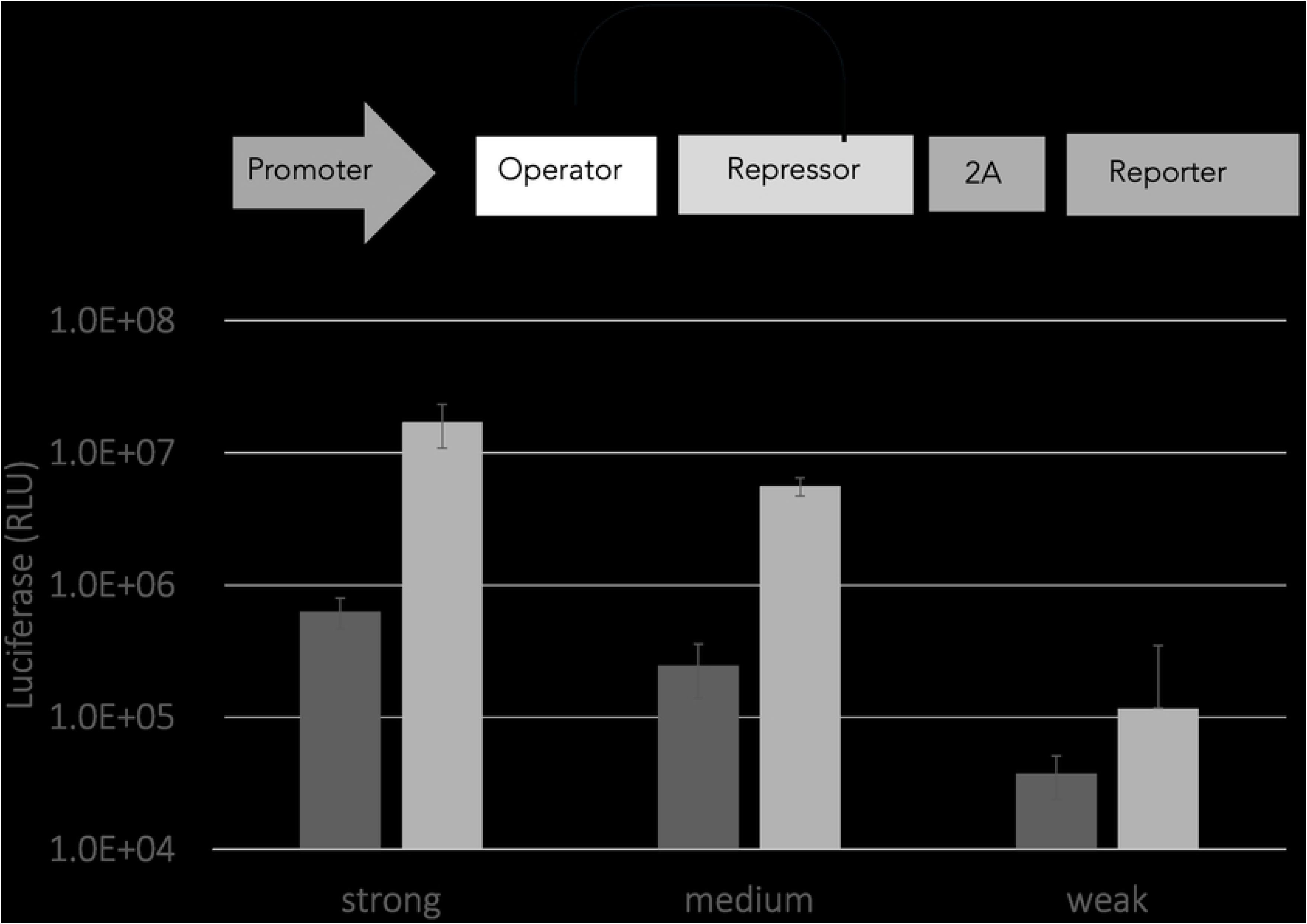
Autogenous TetR regulation across promoters of varying intrinsic strength. A single promoter drives expression of both the luciferase reporter and *cis*-encoded TetR, with repression mediated through an embedded operator sequence. Reporter activity is shown in the absence (OFF) or presence (ON) of doxycycline for weak, medium, and strong promoters. Dark bars denote basal expression; light bars denote induced expression. Error bars represent mean ± SD (*n* = 6). Increasing promoter strength elevates overall expression, whereas TetR-mediated negative feedback suppresses basal output and preserves a relatively constant dynamic range across promoter strengths.

In contrast to the bidirectional architecture, the autogenous system displayed a buffered response to changes in promoter strength. Increasing promoter strength elevated both basal and induced expression, but these increases were sublinear relative to the change in promoter activity. Sublinear scaling, in which output increases less than proportionally with input, is a hallmark of negative feedback regulation and reflects the intrinsic constraints imposed by autoregulatory circuits [10, 12, 14, 20]. Strong promoters generated higher overall expression, but this increase was partially offset by the concurrent rise in TetR production, which feeds back to suppress transcription.

As a result, basal and induced expression varied over a substantially narrower range than in the bidirectional architecture. Although absolute expression levels increased with promoter strength, basal and induced states remained strongly correlated and scaled together across the promoter series. Consequently, fold induction remained relatively stable over much of the promoter range despite large differences in intrinsic promoter activity. These behaviors reflect the fact that promoter strength acts primarily as a global scaling parameter, whereas negative feedback constrains the magnitude of the response and buffers the system against large fluctuations in expression.

Together, these results demonstrate that autogenous regulation fundamentally alters the relationship between promoter strength and gene expression. By embedding negative feedback directly within the regulatory circuit, the architecture suppresses leakiness, reduces expression variability, and stabilizes dynamic range. However, these advantages are accompanied by constraints on maximal output, revealing an intrinsic trade-off between robustness and expression capacity in feedback-regulated systems.

### Thermodynamic model captures autogenous transgene expression

To understand how negative feedback shapes gene expression in the autogenous architecture, we developed a simple thermodynamic model in which transcription depends on promoter occupancy [8–10]. In this framework, the promoter can exist in three mutually exclusive states: unbound, occupied by RNA polymerase (RNAP), or bound by repressor. These states are assigned statistical weights of 1, *α*, and *ρ*(*L*), respectively, where *α* represents effective promoter strength and *ρ*(*L*) describes ligand-dependent repressor occupancy.

Because repressor binding prevents RNAP binding, gene expression is proportional to the probability that RNAP occupies the promoter,

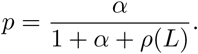

The defining feature of the autogenous circuit is that the repressor is produced from the same promoter that it regulates. Repressor abundance therefore increases automatically as transcription increases, creating negative feedback. To capture this behavior, we assume that repressor occupancy scales with promoter activity,

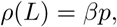

where *β* is an effective feedback parameter that reflects repressor production, stability, and DNA-binding affinity.

Substituting this relation into the promoter occupancy equation gives

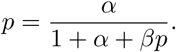

This expression describes a self-consistent feedback loop in which increased transcription produces additional repressor, which in turn suppresses further transcription. Solving for *p* yields

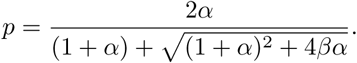

In the absence of feedback (*β* = 0), the model reduces to the standard promoter occupancy relation,

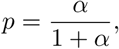

in which expression increases with promoter strength and eventually approaches saturation. In contrast, negative feedback suppresses this increase by coupling higher transcriptional activity to higher repressor production. Under strong-feedback conditions, expression increases only sublinearly with promoter strength,

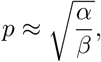

demonstrating that feedback buffers the system against large changes in promoter activity.

The model captures the principal trends observed experimentally in the autogenous architecture. Both basal and induced expression increase with promoter strength, but the increase is coordinated and sublinear, consistent with feedback-mediated buffering. Medium and strong promoters therefore exhibit similar fold induction despite substantial differences in intrinsic transcriptional capacity.

The weakest promoter deviated somewhat from the idealized model, showing reduced induction and lower overall expression. This behavior likely reflects effects not included in the equilibrium framework, such as insufficient repressor accumulation, transcriptional bursting, or resource limitations at very low expression levels. Nevertheless, the overall agreement demonstrates that a minimal self-consistent thermodynamic model is sufficient to explain the major scaling behavior of the autogenous circuit without requiring promoter-specific parameter adjustments.

The model also clarifies the relationship between basal and induced states. In the absence of doxycycline, TetR occupancy is high, corresponding to stronger effective feedback and lower basal expression. Addition of doxycycline reduces TetR DNA-binding affinity, weakening feedback and permitting higher transcriptional output. However, because both OFF and ON states are governed by the same feedback relationship, increases in promoter strength shift both states upward together rather than independently.

Thus, autogenous regulation suppresses leakiness and stabilizes fold induction, but does not completely decouple basal and induced expression. Residual basal expression therefore remains detectable, particularly at higher promoter strengths where increased transcriptional capacity elevates both OFF and ON states. These observations motivated the addition of a second regulatory layer operating at the RNA level. We therefore examined whether ligand-responsive aptazymes could further suppress basal expression while preserving inducibility.

### Dual-layer regulation can suppress leakiness

Regulatory performance can be further improved by combining transcriptional and post-transcriptional control. Ligand-responsive aptazymes provide a natural complement to transcriptional repression by coupling small-molecule binding to mRNA stability through ligand-dependent ribozyme cleavage [15–17]. In the absence of ligand, the ribozyme self-cleaves, destabilizing the transcript and reducing gene expression. Ligand binding inhibits cleavage, stabilizing the mRNA and restoring expression. Although aptazymes alone generally provide only modest regulation, they can further suppress basal expression when combined with transcriptional control [15, 17].

To test this dual-layer strategy, we incorporated the tetracycline-responsive hammerhead aptazymes K7 and K19 into autogenous TetR circuits by inserting the ribozyme sequence into the 3*^′^* untranslated region of the luciferase reporter. These constructs, designated CMV–TetR–K7 and CMV–TetR–K19, combine TetR-mediated transcriptional repression with ligand-responsive RNA destabilization (Fig. 3). Standalone aptazyme reporters (CMV–K7 and CMV–K19) were also generated for comparison.

**Fig 3.**
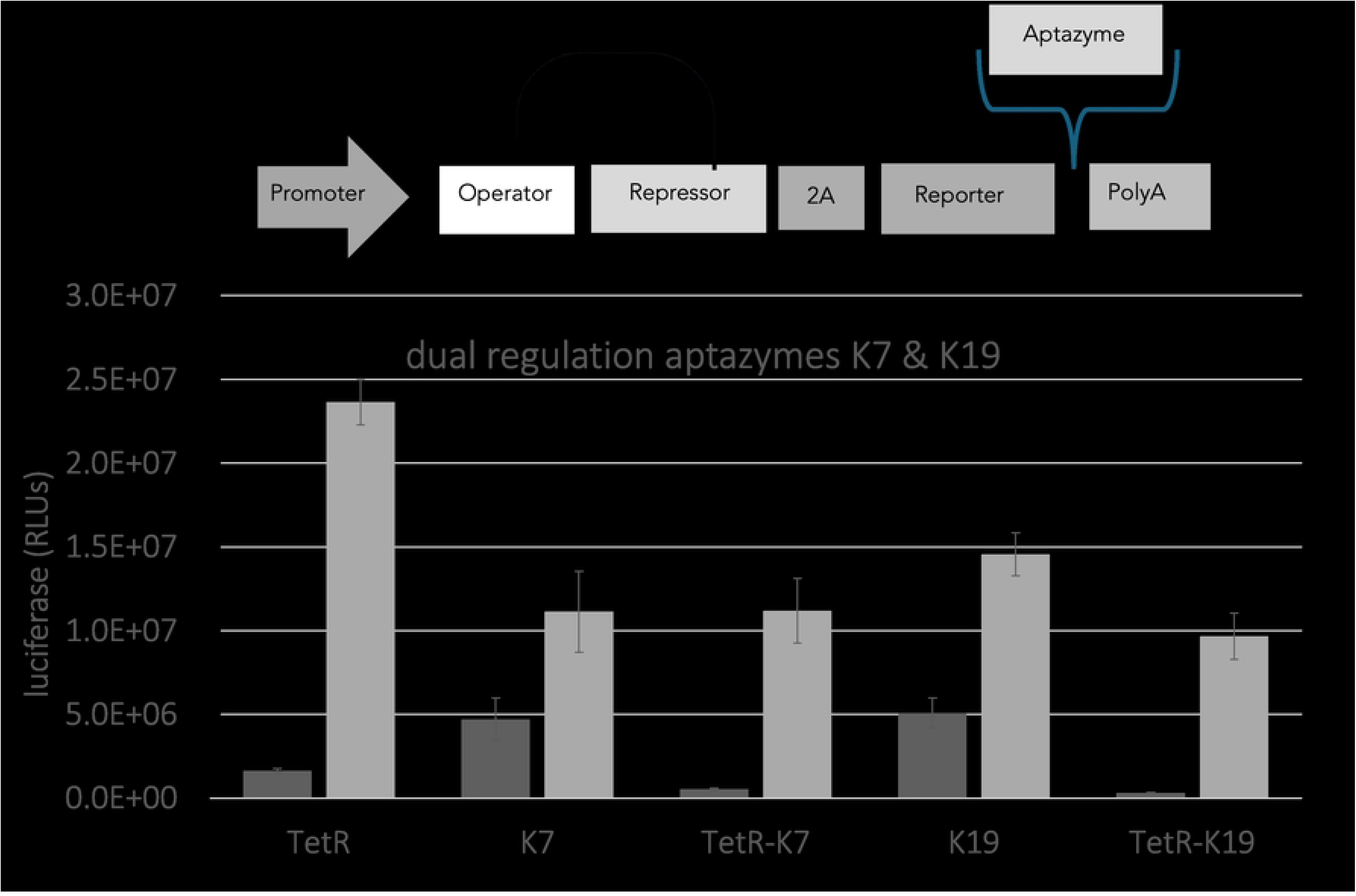
Dual-layer control of TetR-regulated expression using ligand-responsive aptazymes K7 and K19. Reporter output is shown for TetR alone or in combination with aptazymes. Dark bars denote basal (OFF) expression; light bars denote induced (ON) expression. Error bars represent mean ± SD (*n* = 6). Aptazyme incorporation reduces basal expression, whereas combination with TetR further suppresses background while preserving inducibility, demonstrating synergistic transcriptional and post-transcriptional regulation.

Aptazymes alone failed to provide stringent regulation, exhibiting elevated basal expression and limited induction relative to TetR-based repression (Fig. 3), consistent with previous studies of ligand-responsive ribozyme switches in mammalian systems [16, 17]. K7 and K19 alone produced only approximately 2.3-fold and 3-fold induction, respectively. By comparison, TetR alone generated roughly 15-fold induction through partial suppression of basal transcription.

Combining aptazymes with TetR substantially improved OFF-state suppression. Relative to TetR alone, the TetR–K7 and TetR–K19 constructs reduced basal expression several-fold, demonstrating that transcriptional and post-transcriptional mechanisms act synergistically to suppress leakiness. Induced expression was modestly reduced in the dual-layer systems, consistent with incomplete inhibition of ribozyme cleavage under inducing conditions and a corresponding limitation on maximal transcript stability. Similar reductions were observed in the standalone aptazyme constructs, indicating that this effect is intrinsic to the aptazyme mechanism rather than dependent on circuit architecture.

Despite this reduction in maximal output, the dual-layer systems displayed substantially improved dynamic range because suppression of basal expression was much greater than the decrease in induced expression. The TetR–K7 construct exhibited approximately 22-fold induction, whereas TetR–K19 achieved nearly 40-fold induction, representing the strongest overall regulation observed in the study. TetR–K19 reduced basal expression nearly an order of magnitude relative to TetR alone while maintaining induced expression within the same general range.

Together, these results demonstrate that combining transcriptional and RNA-level regulation can markedly improve regulatory stringency by preferentially suppressing OFF-state leakiness. More broadly, the data illustrate a hierarchy of regulatory control in which additional regulatory layers progressively reduce basal expression while also imposing modest constraints on maximal output. Promoter architecture defines the overall relationship between basal and induced expression, whereas post-transcriptional regulation provides an additional layer for tightening OFF-state repression.

## Discussion

The central conclusion of this study is that regulatory architecture sets the operating rules of inducible gene circuits. By combining quantitative experiments with thermodynamic modeling, we show that promoter organization alone is sufficient to generate distinct and predictable behaviors in bidirectional, coupled-promoter, autogenous, and dual-layer regulatory systems. These differences do not arise from changes in repressor chemistry or ligand affinity, but from how promoter strength, repressor abundance, feedback, and regulatory layers are arranged within the circuit. Architecture therefore determines the balance between basal expression, dynamic range, robustness, and maximal output, consistent with general principles of gene network organization [14, 18].

In bidirectional architectures, repressor abundance is controlled by a separate constitutive promoter and is therefore decoupled from the regulated promoter. Although this design enables straightforward tuning of expression strength, it intrinsically links OFF-state leakiness to ON-state output. As promoter strength increases, basal and induced expression rise together, compressing fold induction. These behaviors follow directly from equilibrium occupancy models in which repression and induction reflect the same underlying balance between operator-bound and transcriptionally active promoter states [8–10, 19]. Consequently, bidirectional systems become increasingly sensitive to promoter strength and cellular context at high transcriptional capacity.

These limitations become more pronounced when neighboring promoters are transcriptionally coupled. Incorporating cooperative RNA polymerase–RNA polymerase interactions explains the observed increase in both basal and induced expression that cannot be accounted for by repressor abundance alone. In this regime, stronger promoters stabilize transcriptionally active states, increasing output while simultaneously elevating background expression. Promoter coupling therefore violates the assumption of promoter independence and illustrates how compact genetic arrangements can introduce hidden interactions that dominate circuit behavior [11]. The trade-off is therefore explicit: coupling enhances expression strength at the expense of regulatory precision.

Autogenous regulation provides a distinct solution to these limitations. When the repressor is expressed from the same promoter that it regulates, repressor abundance becomes a self-consistent state variable rather than an externally imposed parameter. Negative feedback buffers promoter-dependent variation and constrains the system to stable operating regimes in which basal and induced expression scale sublinearly with promoter strength while fold induction remains comparatively stable. Such behavior is a hallmark of autoregulatory networks and is associated with reduced noise and improved robustness [12–14, 20]. Our thermodynamic framework captures these effects without invoking additional regulatory complexity, demonstrating that feedback alone reshapes the equilibrium landscape governing repression and induction.

The aptazyme experiments further demonstrate that additional regulatory layers can improve control when transcriptional repression alone is insufficient. Aptazymes alone displayed limited dynamic range because elevated basal expression constrained induction. In contrast, combining transcriptional repression with RNA-level control produced substantially larger fold induction by selectively suppressing basal expression. The TetR–K7 and TetR–K19 circuits exhibited markedly improved dynamic range relative to either component alone, with TetR–K19 providing the strongest OFF-state suppression. This enhanced regulatory precision arose primarily through reduction of basal expression rather than increased maximal output, as induced expression was modestly reduced relative to TetR alone. More generally, different regulatory layers act on distinct aspects of circuit behavior: promoter architecture defines the relationship between basal and induced states, whereas post-transcriptional regulation preferentially suppresses residual leakiness.

Together, these results support a hierarchical view of regulatory design. Independent promoters establish the baseline logic of repression, promoter coupling introduces hidden transcriptional interactions, autogenous feedback stabilizes circuit behavior, and post-transcriptional regulation further suppresses residual basal expression. Although chromatin structure, transcriptional bursting, and other non-equilibrium effects may contribute to quantitative differences, these factors modulate rather than override the underlying thermodynamic constraints imposed by circuit architecture.

More broadly, this work demonstrates that the behavior of inducible gene circuits is determined not only by ligand binding and allostery, but also by how regulatory components are physically organized within the network. Architecture defines the thermodynamic operating regime through which molecular interactions are translated into cellular output. These principles provide a general framework for designing compact, robust, and predictable gene-control systems in mammalian cells.

## Data availability

All data generated or analyzed during this study are reported in the manuscript and its figure files. Raw data is available: https://doi.org/10.5061/dryad.9kd51c606. Plasmid sequences could be made available upon request to the corresponding author.

## Author Contributions

Conceptualization: Mitchell Lewis.

Methodology: Mitchell Lewis, Abhilasha Gupta.

Investigation: Abhilasha Gupta.

Formal analysis: Mitchell Lewis, Abhilasha Gupta.

Writing – original draft: Mitchell Lewis.

Writing – review & editing: Mitchell Lewis, Abhilasha Gupta.

Funding acquisition: Mitchell Lewis.

Supervision: Mitchell Lewis.

## Materials and methods

### Promoter characterization

Promoter activities were measured by transient transfection followed by firefly luciferase assay. The panel comprised CMV, CAG, CMV–IE, long and core EF1*α* variants, CMV_min_, and hPGK. Activity spanned nearly two orders of magnitude, with CMV producing approximately 100-fold more signal than hPGK. CMV, CAG, and CMV–IE formed a high-activity group (approximately 50–100× hPGK), the EF1*α* variants occupied an intermediate range (approximately 8–30× hPGK), and CMV_min_ and hPGK defined the low-activity range. These experimentally determined activities were used to order promoter strength in subsequent comparisons.

### Plasmid construction

All regulatory modules were derived from the AAV-compatible pSW2.Luc backbone, a previously described autogenously regulated expression system flanked by AAV2 inverted terminal repeats (ITRs) [23]. Functional elements in this backbone were bounded by restriction sites to permit modular exchange of promoters, operators, repressors, and post-transcriptional regulatory elements. Synthetic double-stranded DNA fragments (gBlocks; Integrated DNA Technologies) were used for replacement and were cloned directionally with the indicated flanking restriction sites. All final plasmids were verified by bidirectional Sanger sequencing.

The parental autogenous cassette contained, from the 5*^′^*ITR, a CMV immediate-early enhancer/promoter, an operator positioned between the TATA box and transcription start site, a synthetic intron (Promega), the repressor coding sequence, a P2A ribosomal-skipping sequence, the firefly luciferase reporter Luc2 (Promega), an SV40 polyadenylation signal, and the 3*^′^* ITR. TetR-regulated autogenous constructs were generated by replacing the SacI–StuI region containing the lac operators with two copies of *tetO* and replacing the StuI–SpeI-flanked *lacI* coding region with *tetR*. The P2A sequence permitted coordinated expression of TetR and luciferase as separate protein products [24].

Operator number, operator spacing, and the distance between the operator and TATA box were varied by replacing the SacI–StuI region. Promoter variants and truncated CMV derivatives were introduced using NheI–SacI, NdeI–SacI, or SacI–StuI sites, depending on the promoter fragment. K7 and K19 aptazymes, together with the corresponding ribozyme controls, were inserted into the linker between Luc2 and the SV40 polyadenylation signal using NotI and XhoI sites, thereby placing the aptazyme in the reporter 3*^′^* untranslated region.

For bidirectional constructs, the TetR and reporter transcriptional units were placed in divergent orientations and separated by a transcriptional pause sequence. The reverse-oriented promoter drove TetR expression, whereas the opposing promoter–*tetO* cassette drove Luc2. The TetR bidirectional module was generated by replacing the PurR–VP16 coding region of the precursor bidirectional vector with *tetR* and replacing the PurR-responsive promoter/operator region with the corresponding CMV–*tetO* cassette. Reporter-side CMV variants included CMV enhancer plus CMV_min_, CMV–IE, and CMV_min_ constructs with or without operators. Repressor-side hPGK was replaced where indicated with CMV–IE using HindIII and NheI. The resulting promoter combinations were used to vary TetR abundance and reporter transcriptional capacity independently.

All constructs were maintained below 4,274 bp from the 5*^′^*to 3*^′^* ITR, within the approximately 4.7-kb AAV genome packaging capacity [22] (Fig. 4).

**Fig 4.**
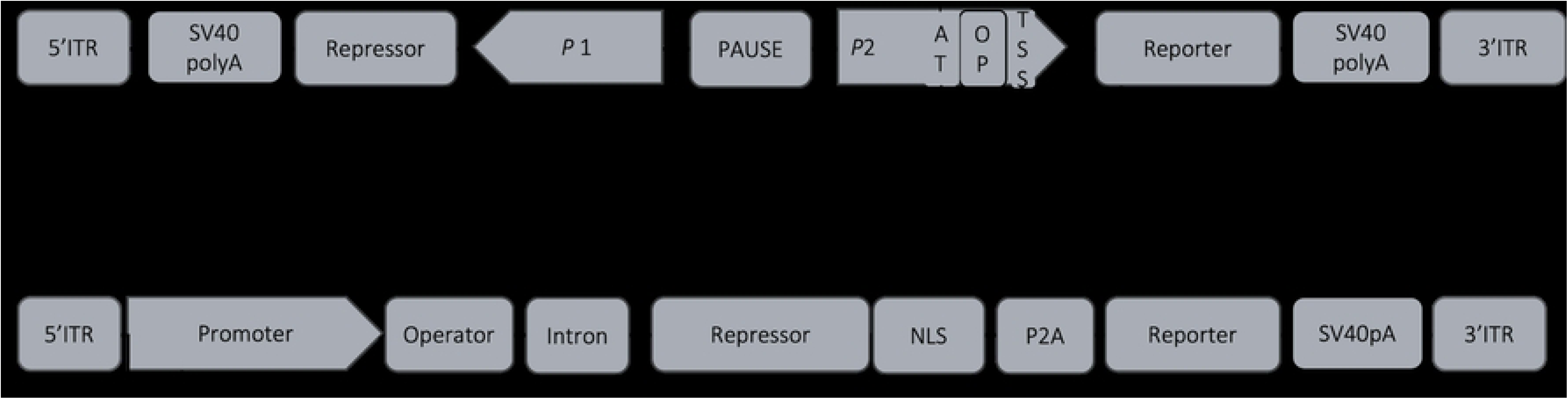
Schematic of bidirectional and autogenous regulatory constructs. Promoter variants and regulatory elements were introduced using standard cloning strategies. All constructs were sequence verified and flanked by AAV2 ITRs.

### Cell culture and transfection

HEK293T (ATCC) were maintained in high-glucose DMEM containing GlutaMAX (Gibco), 10% fetal bovine serum (USDA-tested HyClone; Fisher Scientific), and 1% penicillin/streptomycin (Gibco). Cells were cultured at 37 °C in a humidified atmosphere containing 5% CO_2_.

For plasmid assays, cells were counted with a hemocytometer and seeded in 96-well plates 18 h before transfection. HEK293T cells were seeded at 4 × 10^4^ cells in 100 µL per well in 100 µL per well, yielding approximately 70–90% confluence at transfection. Each well received 20 ng plasmid DNA in a final volume of 110 µL. HEK293T and HeLa cells were transfected with Lipofectamine 2000, whereas ARPE-19 cells were transfected with Lipofectamine LTX plus PLUS Reagent (Thermo Fisher Scientific), following the manufacturer’s instructions. Within each experiment, equal amounts of plasmid DNA were delivered per condition and samples were processed in parallel.

### Induction and expression measurements

Four hours after transfection, TetR-containing cultures were treated with tetracycline or doxycycline, as appropriate for the construct, in a dose-response format and incubated for an additional 20 h at 37 °C. Untreated wells defined basal/OFF expression, and ligand-treated wells defined induced/ON expression. For the comparisons reported here, induced values were taken at the saturating ligand condition used in the corresponding experiment. Aptazyme-containing constructs received tetracycline in addition to the inducer used for the transcriptional module, as applicable.

Firefly luciferase activity was measured with Bright-Glo Luciferase Assay System (Promega). Culture medium was removed and 100 µL Bright-Glo reagent, diluted 1:1 with dye-free RPMI, was added to each well. Luminescence was measured with an Infinite 200 plate reader using Tecan software and reported as relative light units (RLU). Raw signals were background-subtracted before analysis.

### Data analysis

Basal and induced expression levels were defined as

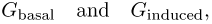

corresponding to reporter expression in the absence and presence of doxycycline, respectively. Fold induction was calculated as

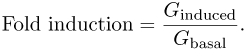

Expression levels were analyzed as a function of promoter strength. For bidirectional architectures, expression was evaluated with respect to both repressor-promoter and reporter-promoter strengths. For autogenous architectures, expression was analyzed as a function of the shared promoter.

Scaling relationships were assessed by comparing basal and induced expression across promoter strengths on linear and logarithmic scales. Correlations between basal and induced states were evaluated by plotting *G*_induced_ versus *G*_basal_.

### Thermodynamic modeling

Regulatory behavior was interpreted using an equilibrium statistical thermodynamic framework in which gene expression is proportional to the probability of RNA polymerase occupancy. Promoter states were enumerated explicitly, and transcriptional output was calculated from the corresponding partition function. Ligand binding modulates the effective repressor concentration by shifting the equilibrium between DNA-binding–competent and incompetent states.

Differences in expression between architectures arise from how promoter occupancy, repressor binding, and promoter–promoter interactions are incorporated into the model.

### Statistical analysis

Each construct and ligand condition was measured in replicate, with the replicate number reported in the corresponding figure legend. Values are shown as mean ± standard deviation. Absolute expression is reported as background-subtracted RLU. Where fractional expression was used, values were normalized to the corresponding constitutive promoter control. Fold induction was calculated as the ratio of maximally induced to basal luciferase activity. No additional normalization was applied unless stated in the figure legend.

